# Interneuronal correlations dynamically adjust to task demands at multiple time-scales

**DOI:** 10.1101/547802

**Authors:** S Ben Hadj Hassen, C. Gaillard, E. Astrand, C. Wardak, S Ben Hamed

**Affiliations:** Institut des Sciences Cognitives Marc Jeannerod, CNRS, UMR5229, 67 Boulevard Pinel, 69675 Bron Cedex, France; Mälardalen University, IDT, Högskoleplan 1, 721 23 Västerås, Sweden; Imagerie et Cerveau (iBrain), 10 Boulevard Tonnellé 37032 Tours Cedex 1, France

**Keywords:** noise correlation, prefrontal cortex, macaque monkey, task difficulty, alpha oscillations, beta oscillations, cognitive demand, cognitive flexibility, rhythmic cognition

## Abstract

Functional neuronal correlations between pairs of neurons are thought to play an important role in neuronal information processing and optimal neuronal computations during attention, perception, decision-making and learning. Here, we report dynamic changes in prefrontal neuronal noise correlations at multiple time-scales, as a function of task contingencies. Specifically, we record neuronal activity from the macaque frontal eye fields, a cortical region at the source of spatial attention top-down control, while the animals are engaged in tasks of varying cognitive demands. We show that the higher the task demand and cognitive engagement the lower noise correlations. We further report that within a given task, noise correlations significantly decrease in epoch of higher response probability. Last we show that the power of the rhythmic modulations of noise correlations in the alpha and beta frequency ranges also decreases in the most demanding tasks. All of these changes in noise correlations are associated with layer specific modulations in spikes-LFP phase coupling, suggesting both a long-range and a local intra-areal origin. Over all, this indicates a highly dynamic adjustment of noise correlations to ongoing task requirements and suggests a strong functional role of noise correlations in cognitive flexibility.

**Significance statement:** Cortical neurons are densely interconnected. As a result, pairs of neurons share some degree of variability in their neuronal responses. This impacts how much information is present within a neuronal population and is critical to attention, decision-making and learning. Here we show that, in the prefrontal cortex, this shared inter-neuronal variability is highly flexible, decreasing across tasks as cognitive demands increase and within trials in epochs of maximal behavioral demand. It also fluctuates in time at a specific rhythm, the power of which decreases for higher cognitive demand. All of these changes in noise correlations are associated with layer specific modulations in spikes-LFP phase coupling. Over all, this suggests a strong functional role of noise correlations in cognitive flexibility.

## Introduction

Optimal behavior is the result of interactions between neurons both within and across brain areas. Identifying how these neuronal interactions flexibly adjust to the ongoing behavioral demand is key to understand the neuronal processes and computations underlying optimal behavior. Several studies have demonstrated that functional neuronal correlations between pairs of neurons, otherwise known as noise correlations, play an important role in perception and decision-making^1–9^. Specifically, several experimental and theoretical studies show that noise correlations have an impact on the amount of information that can be decoded for neuronal populations^4, 10–12^ as well as on overt behavioral performance^4, 10–15^. As a result, understanding how noise correlations dynamically adjust to task demands is a key step toward clarifying how neural circuits dynamically control information transfer, thereby optimizing behavioral performance.

Several sources of noise correlations have been proposed, ranging from shared connectivity^16^, to global fluctuations in the excitability of cortical circuits^17, 18^, feedback signals ^19^ or internal areal dynamics^20–22^, or bottom-up peripheral sensory processing^23^. In an independent study ^24^, we for example show that noise correlations in the prefrontal cortex fluctuate rhythmically in the high alpha (10-16Hz) and beta (20-30Hz) frequency ranges. We further show that these rhythmic fluctuations co-occur with changes in spike-LFP phase coupling and can be segregated into a long-range component, possibly reflecting global fluctuations in the excitability of the functional network of interest, as well as to local internal areal dynamics. Importantly, these fluctuations account for variability in behavioral performance.

From a cognitive point of view, noise correlations have been shown to change as a function of spatial attention^25^, spatial memory ^26^ and learning^27, 28^, suggesting that they are subject both to rapid dynamic changes as well as to longer term changes, supporting optimal neuronal computations^28^.

Here, we focus onto how the strength of rhythmic modulation of noise correlations, in the prefrontal cortex, is affected by the ongoing task at multiple time-scales. Specifically, we record neuronal activity from the macaque frontal eye fields, a cortical region which has been shown to be at the source of spatial attention top-down control ^15, 29–31^ while the animals are engaged in tasks of varying cognitive demands, as assessed by their overt behavioral performance. Overall, we show that noise correlations vary in strength both as a function of the ongoing task as well as a function of the probabilistic structure of the task, thus adjusting dynamically to ongoing behavioral demands. Importantly, we show that these variations of noise correlations strength across tasks co-occur with cortical layer specific variations in spikes-LFP phase coupling.

## Results

Our main goal in this work is to examine how the degree of cognitive engagement and task demands impact the neuronal population state as assessed from inter-neuronal noise correlations. Cognitive engagement was operationalized through tasks of increasing behavioral requirements. The easiest task (***Fixation task***, figure 1B.1) is a central fixation task in which monkeys are required to detect an unpredictable change in the color of the fixation point, by producing a manual response within 150 to 800ms from color change. The second task (***Target detection task***, figure 1B.2) adds a spatial uncertainty on top of the temporal uncertainty of the event associated with the monkeys’ response. It is a target detection task, in which the target can appear at one of four possible locations, at an unpredictable time from fixation onset. The monkeys have to respond to this target presentation by producing a manual response within 150 to 800ms from color change. In the third task (***Memory guided saccade task***, figure 1B.3), monkeys are required to hold the position of a spatial cue in memory for 700 to 1900ms and to perform a saccade towards that memorized spatial location on the presentation of a go signal. This latter task thus involves a temporal uncertainty but no spatial uncertainty. However, in contrast with the previous tasks, it requires the production of a spatially oriented oculomotor response rather than a simple manual response. Accordingly, both monkeys have higher performances on the memory guided saccade task than on the target detection task (Figure 1C, Wilcoxon rank sum test, Monkey 1, p<0.01, Monkey 2, p<0.05), and higher performances on the target detection task than on the fixation task (Wilcoxon rank sum test, p<0.05). Importantly, task order was randomized from one session to the next, such that the reported effects could not be accounted for by fatigue or satiation effects.

**Figure 1:**
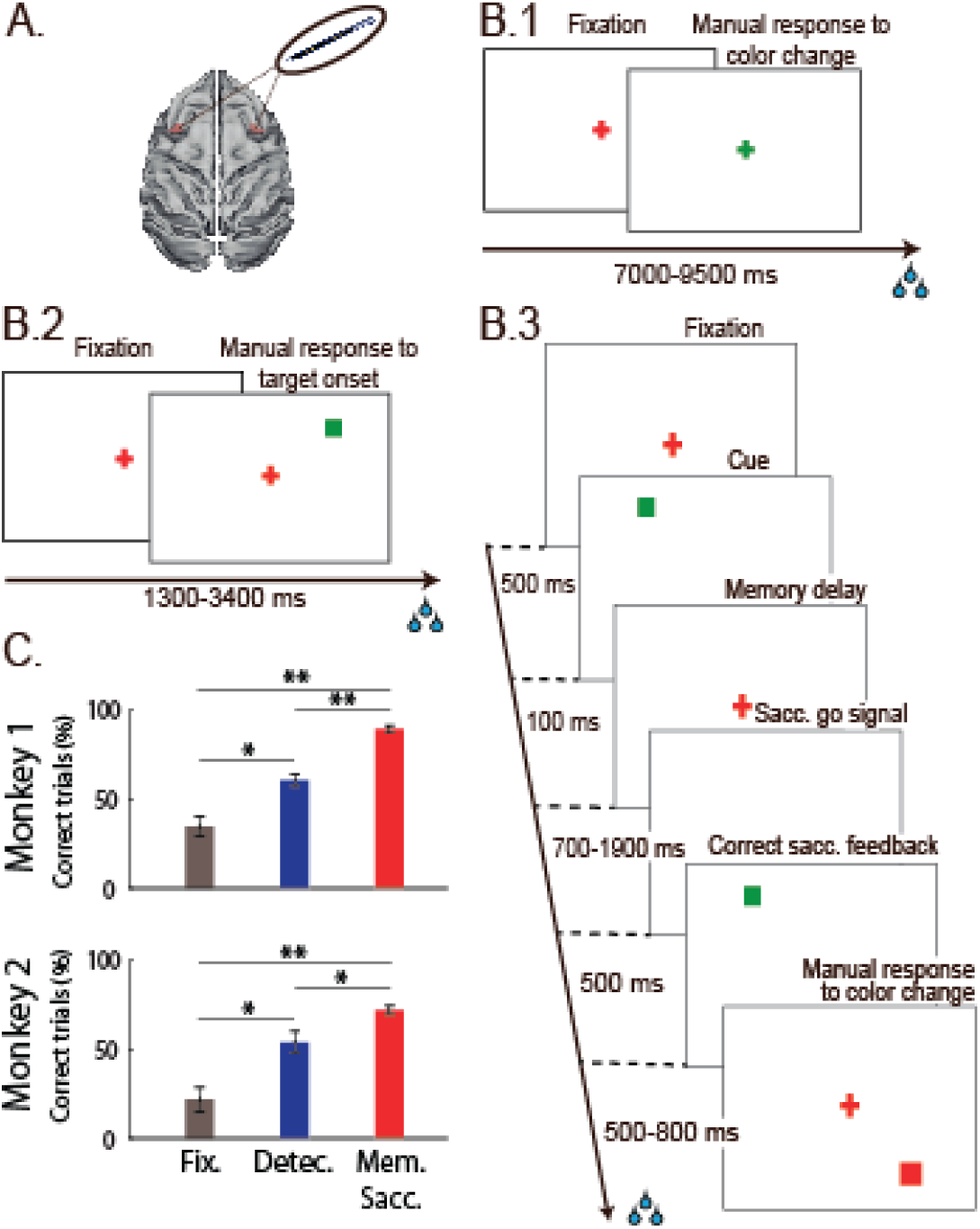
(A*) **Recordings sites.** On* each session, 24-contacts recording probes were placed in the left and right FEFs. (B.1) ***Fixation task.*** Monkeys had to fixate a red central cross and were rewarded for producing a manual response 150ms to 800ms following fixation cross color change. (B.2) ***Target detection task.*** Monkeys had to fixate a red central cross and were rewarded for producing a manual response 150ms to 800ms from the onset of a low luminosity target at an unpredictable location out of four possible locations on the screen. (B.3) ***Memory-guided saccade task.*** Monkeys had to fixate a red central cross. A visual cue was briefly flashed in one of four possible locations on the screen. Monkeys were required to hold fixation until the fixation cross disappeared and then produce a saccade to the spatial location indicated by the cue within 300ms from fixation point offset. On success, the cue re-appeared and the monkeys had to fixate it. They were then rewarded for producing a manual response 150ms to 800ms following the color change of this new fixation stimulus. (C) ***Behavioral performance.*** Average percentage of correct trials across sessions for each tasks and each monkey with associated standard errors.

Neuronal recordings were performed in the prefrontal cortex, specifically in the frontal eye field (FEF, figure 1A), a structure known to play a key role in covert spatial attention^31–34^. In each session, multi-unit activity (MUA) and local field potential (LFP) were recorded bilaterally, while monkeys performed these three tasks. In the following, noise correlations between the different prefrontal signals of the same hemisphere were computed on equivalent task fixation epochs, away from both sensory intervening events and motor responses. We analyzed how these noise correlations varied both across tasks, as a function of cognitive engagement and within-tasks, as a function of the probabilistic structure of the task.

### Noise correlations decrease as cognitive engagement and task requirements increase

In order to characterize how inter-neuronal noise correlations vary as a function of cognitive engagement and task requirements, we proceeded as follows. In each session (n=26), noise correlations were computed between each pair of task-responsive channels (n=671, see Methods), over equivalent fixation task epochs, running from 300 to 500 ms after eye fixation onset. This epoch was at a distance from a possible visual or saccadic foveation response and in all three tasks, monkeys were requested to maintain fixation at this stage. It was also still early on in the trial, such that no intervening sensory event was to be expected by the monkey at this time. Importantly, fixation behavior, i.e. the distribution of eye position in within the fixation window, did not vary between the different tasks (Friedman test, p<0.001). As a result, and because tasks were presented in blocks, any difference in noise correlations across tasks during this “neutral” fixation epoch are to be attributed to general non-specific task effects, i.e. differences in the degree of cognitive engagement and task demands. Noise correlations were significantly different between tasks (figure 2, ANOVA, p<0.001). Specifically, they were higher in the fixation task than in the target detection task (figure 2, Wilcoxon rank sum test, p<0.001) and in the memory guided saccade task (Wilcoxon rank sum test, p<0.001). They were also significantly higher in the target detection task than in the memory guided saccade task (Wilcoxon rank sum test, p<0.001). Importantly, these significant changes in noise correlations existed in the absence of significant differences in mean firing rate (ANOVA, p>0.5), standard error around this mean firing rate (ANOVA, p>0.6), and Fano factor (ANOVA, p>0.7). These task differences in noise correlations were preserved as noise correlations decreased as a function on the cortical distance between the neuronal pairs (figure S1A). These task differences were also preserved across pairs sharing the spatial functional selectivity or not (figure S1B). We thus describe that, in absence of any sensory or cognitive processing, noise correlations are strongly modulated by cognitive engagement and task demands.

**Figure 2:**
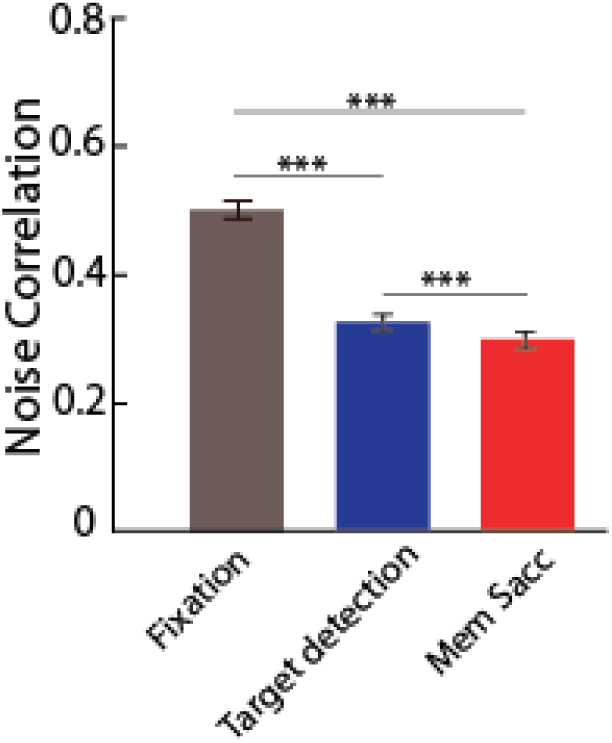
Noise correlations as a function of task. Average noise correlations across sessions for each of the three tasks (mean +/- s.e., noise correlations calculated on the neuronal activities from 300 to 500 after eye fixation onset. Grey: fixation task; blue: target detection task; red: memory guided saccade task. Stars indicate statistical significance following a one-way ANOVA; *p<0.05; **p<0.01; ***p<0.001.

The task differences in noise correlations described above could reflect changes in the shared functional connectivity, within the large-scale parieto-frontal functional network the cortical region of interest belongs to ^16^ or to global fluctuations in the excitability of cortical circuits ^35, 36^. This large-scale hypothesis predicts that the observed changes in noise correlations are independent from intrinsic connectivity as assessed by the distance, the spatial selectivity and the cortical layer between the pairs of signals across which noise correlations are computed. Alternatively, these task differences in noise correlations could reflect a more complex reweighing of functional connectivity and the excitatory/inhibitory balance in the area of interest, due to local changes in the random shared fluctuations in the pre-synaptic activity of cortical neurons ^4, 16, 37, 38^. This local hypothesis predicts that the observed changes in noise correlations depend onto intrinsic microscale connectivity.

FEF neurons are characterized by a strong visual, saccadic, spatial memory and spatial attention selectivity ^31, 39, 40^. Previous studies have shown that pure visual neurons are located in the input layers of the FEF while visuo-motor neurons are located in its output layers ^39, 41–45^. Independently, it has been shown that, in extrastriate area V4, the ratio between the alpha and gamma spike field coherence discriminated between LFP signals in deep (low alpha / gamma spike field coherence ratio) or superficial cortical layers (high alpha / gamma spike field coherence ratio) ^46^. In our own data, because our recordings were performed tangentially to FEF cortical surface, we have no direct way of assigning the recorded MUAs to either superficial or deep cortical layers. However, the alpha / gamma spike field coherence ratio provides a very reliable segregation of visual and viso-motor MUAs (figure 3A). We thus consider that, as has been described for area V4, this measure allows for a robust delineation of superficial and deep layers in area FEF. In the following, we computed inter-neuronal noise correlations between three different categories of pairs based on their assigned cortical layer: superficial/superficial pairs, superficial/deep pairs and deep/deep pairs, where superficial MUA correspond to predominantly visual, low alpha/gamma spike field coherence ratio signals and deep MUA correspond to predominantly visuo-motor, high alpha/gamma spike field coherence ratio signals. Noise correlations varied as a function of cortical layer (Figure 3B). This cortical layer effect was present for all tasks and expressed independently of the main task effect described above (2-way ANOVA, Task x Cortical layer, Task effect: p<0.001; Cortical layer effect: p<0.001). Layer effects were not constant across tasks, possibly suggesting task-dependent functional changes in within and across layer neuronal interactions (interaction: p<0.05). Unexpectedly, belonging to the same layer cortical layer didn’t systematically maximize noise correlations. Indeed, post-hoc analyses indicate significantly lower noise correlations between the superficial/superficial pairs as compared to the deep/deep pairs (Wilcoxon rank sum test, Fixation task: p<0.05; Target detection task: p<0.05; Memory-guided saccade task: p<0.01). Superficial/deep pairs sat in between these two categories and had significantly lower noise correlations than the deep/deep pairs (Wilcoxon rank sum test, Fixation task: p<0.05; Target detection task: p<0.05; Memory-guided saccade task: p<0.01) and higher noise correlations than the superficial/superficial pairs, though this difference was never significant.

**Figure 3:**
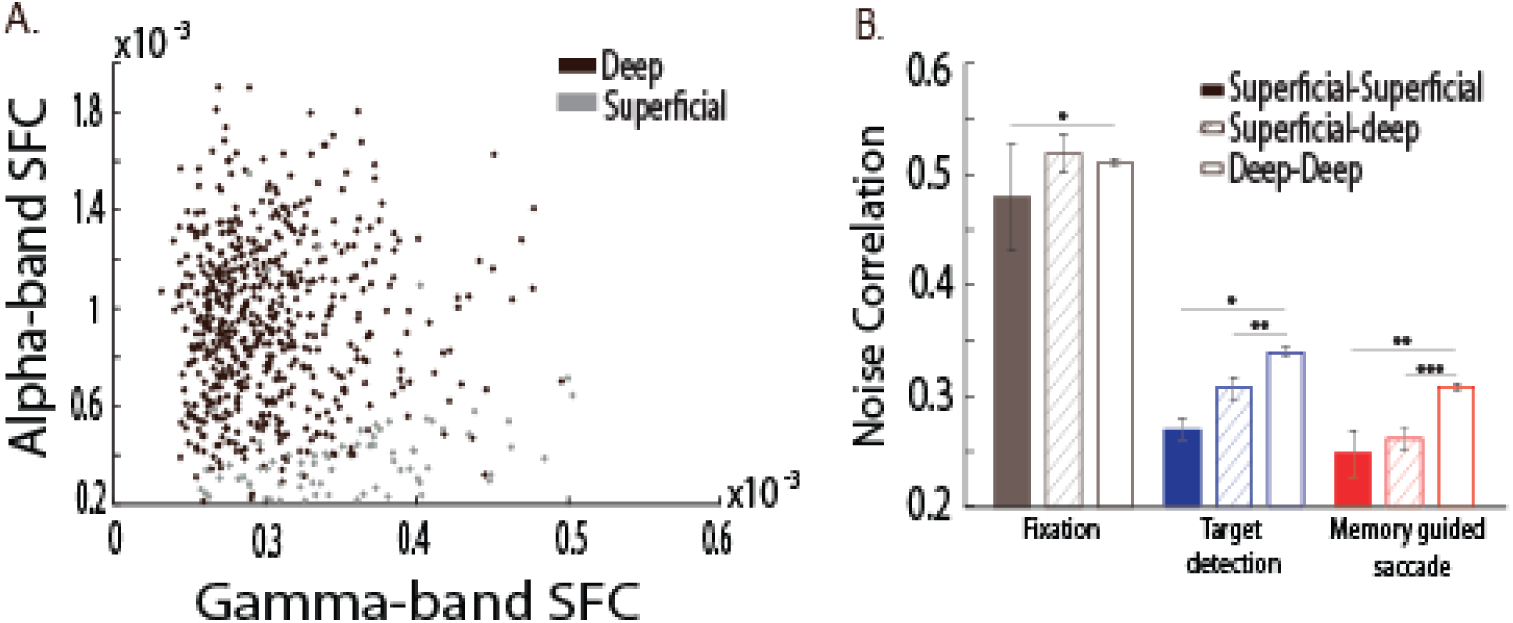
(A) Distribution of alpha spike-LFP coupling (6-16Hz) as a function of gamma (40-60Hz) spike-LFP coupling for visual and visuomotor frontal eye field sites. Sites with visual selectivity but no motor selectivity (grey, putative superficial sites) demonstrated stronger gamma-band spike-LFP phase coupling, whereas sites with visuomotor selectivity (black, putative deep sites) demonstrated stronger alpha-band spike-LFP coupling. (B) Noise correlations as a function of pair functional selectivity. Average of noise correlations (mean +/- s.e.) across sessions, for each task, from 300ms to 500ms after eye fixation onset, as a function of pair functional selectivity: visual-visual, visual-visuomotor, visuomotor-visuomotor. Stars indicate statistical significance following a two-way ANOVA and ranksum post-hoc tests; *p<0.05; **p<0.01; ***p<0.001.

Overall, these observations support the co-existence of both a global large-scale change as well as a local change in functional connectivity. Indeed, task effects onto noise correlations build up onto cortical distance, spatial selectivity and cortical layer effects, indicating global fluctuations in the excitability of cortical circuits ^35, 36^. On top of this global effect, we also note more complex changes as reflected from statistical interactions between task and spatial selectivity or layer attribution effects. This points towards more local changes in neuronal interactions, based on both 1) functional neuronal properties such as spatial selectivity that may change across tasks ^47–50^ and 2) the functional reweighing of top-down and buttom-up processes^29, 31^.

### Impact of the probabilistic structure of the task onto noise correlations

The variation of noise correlations as a function of cognitive engagement and task demands suggests a flexible adaptive mechanism that adjusts noise correlations to the ongoing behavior. On task shifts, this mechanism probably builds up during the early trials of the new task, past trial history affecting noise correlations in the current trials. In a previous study^51^, we show that, in a cued target detection task, while noise correlations are higher on miss trials than on hit trials, noise correlations are also higher on both hit and miss trials, when the previous trial was a miss as compared to when it was a hit. Here, one would expect that on the first trials of task shifts, noise correlations would be at an intermediate level between the previous and the ongoing task. Task shifts being extremely rare events in our experimental protocol, this cannot be confirmed. On top of this slow dynamics carry on effect, one can also expect faster dynamic adjustments to the probabilistic structure of the task. This is what we demonstrate below.

In each of the three tasks, target probability (saccade go signal probability in the case of the memory guided saccade task) varied as a function of time. As a result, early target onset trials had a different target probability compared to intermediate and late target onset trials. Our prediction was that if monkeys had integrated the probabilistic structure of the task, this should reflect onto a dynamic adjustment of noise correlations as a function of target probability. Figure 4 confirms this prediction. Specifically, for all tasks, noise correlations were lowest in task epochs with highest target probability (Wilcoxon non-parametric test, p<0.001 for all pair-wise comparisons). These variations between the highest and lowest target probability epochs were highly significant and in the order of 15% or more (Fixation task: 15%, Target detection task: 40%, Memory-guided saccade task: 14%). This variation range was lower than the general task effect we describe above but yet quite similar across tasks. Overall, this indicates that noise correlations in addition to being dynamically adjusted to task structure, are also dynamically adjusted to trial structure, and are lowest at the time of highest behavioral demand in the trial.

**Figure 4:**
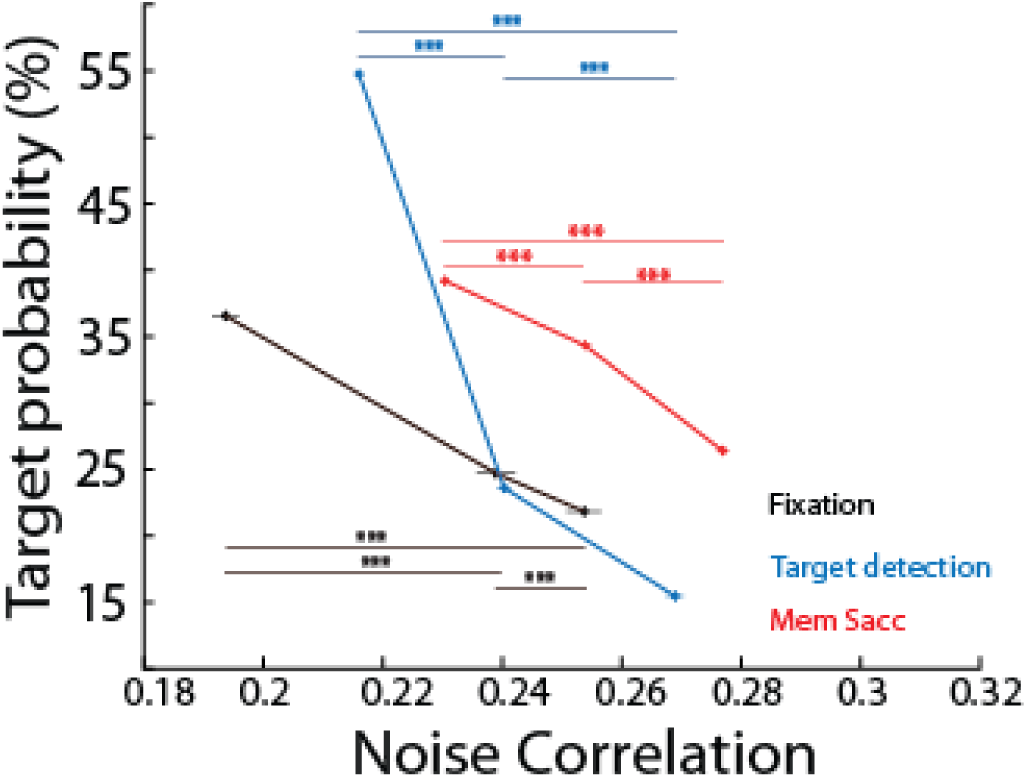
Noise correlations decrease as function of expected response probability. Average noise correlations (mean +/- s.e.) across sessions, for each task, calculated on 200ms before the target (fixation and target detection tasks) onset or saccade execution signal onset (memory guided saccade task), as a function of expected target probability. Each data point corresponds to noise correlations computed over trials of different fixation onset to event response intervals, i.e. over trials of different expected response probability. Stars indicate statistical significance following a two-way ANOVA and ranksum post-hoc tests; *p<0.05; **p<0.01; ***p<0.001.

### Strength of rhythmic fluctuations in noise correlations as a function of tasks

In an independent study ^24^, we show that noise correlations in the prefrontal cortex fluctuate rhythmically in the high alpha (10-16Hz) and beta (20-30Hz) frequency ranges. This is reproduced here in three distinct tasks (figure 5A). Noise correlations phase locked to fixation onset (Fixation and target detection task) or cue presentation (Memory guided saccade task) are characterized by rhythmic fluctuations in two distinct frequency ranges: a high alpha frequency range (10-16 Hz) and a beta frequency range (20-30Hz), as quantified by a wavelet analysis (figure 5B). These oscillations can be described in all of the three tasks, this in spite of an overall higher background spectral power during the memory guided saccade task, both when noise correlations are calculated on trials in which spatial memory was instructed towards the preferred or the non-preferred location of the MUA signals (figure 5B, red and green curves respectively). Because spatial selective processes are at play in the memory guided saccade task, both for trials in which spatial memory is oriented towards the preferred MUA location (excitatory processes) or towards the non-preferred location (inhibitory processes), we will mostly focus on the fixation and the target detection tasks.

**Figure 5:**
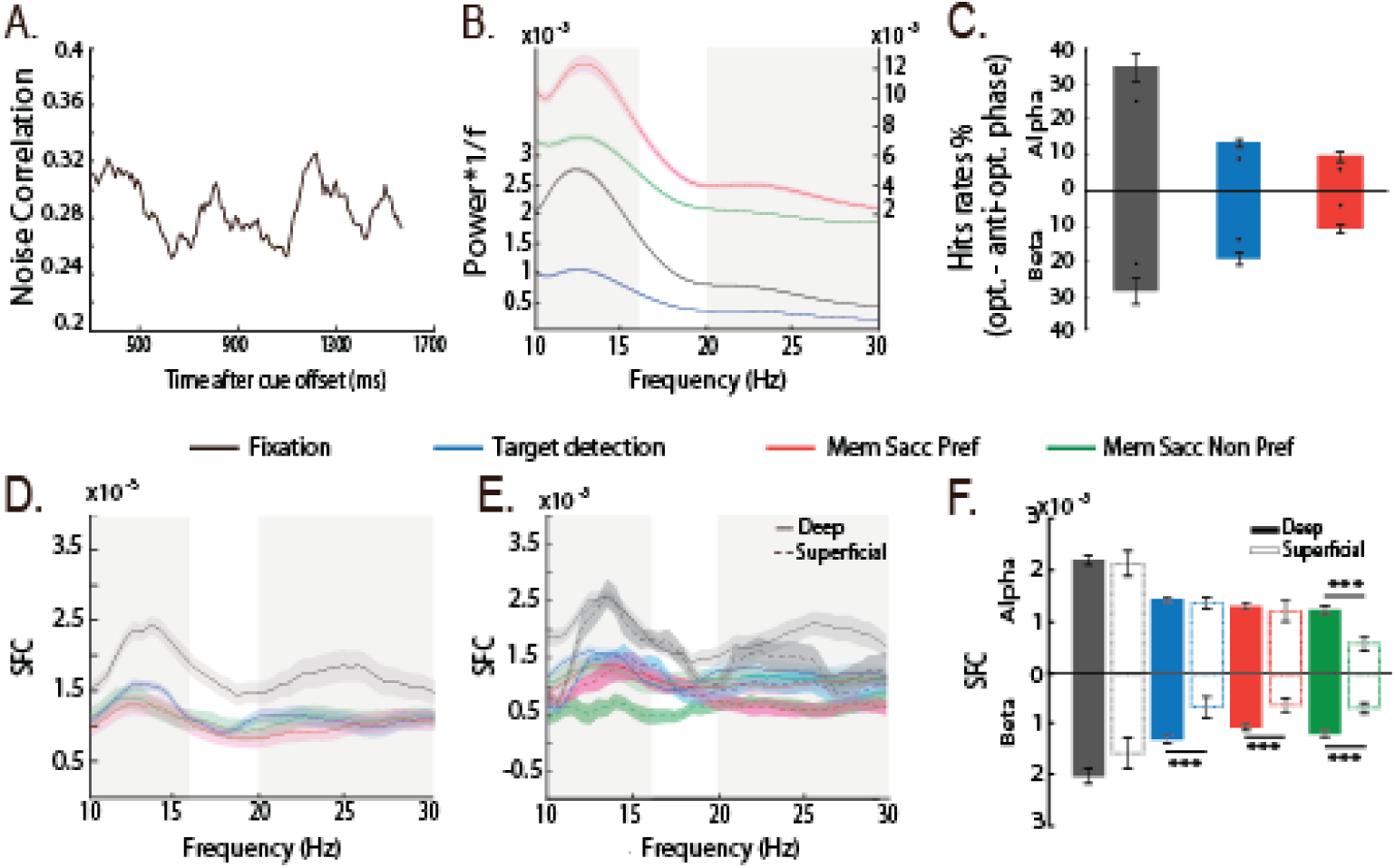
Rhythmic fluctuations in noise correlations modulate behavioral response and spike-LFP phase coupling in upper input cortical layers. (A) Single memory guided saccade session example of noise correlation variations as a function of trial time. (B) 1/f weighted power frequency spectra of noise correlation in time (average +/- s.e.m), for each task, calculated from 300ms to 1500ms from fixation onset (Fixation and Target detection tasks) or following cue offset (Memory guided saccade task). (C) Hit rate modulation by alpha (top histogram) and beta (bottom histogram) noise correlation at optimal phase as compared to anti-optimal phase for all three tasks (color as in (B), average +/- s.e., dots represent the 95% confidence interval under the assumption of absence of behavioral performance phase dependence). (D) Spikes-LFP phase coupling between LFP and spike data as a function of frequency, time intervals as in (B). (E) Spikes-LFP phase coupling calculated as in (C) but as a function of the layer attribution of each signal, time intervals as in (B). (F) Average SFC (+/- s.e.) in alpha (10-16Hz, top histogram) and beta (20-30Hz, bottom histogram) for each task and both of superficial and deep cortical layer signals (t-test, ***: p<0.001).

When compensating the rhythmic modulations of noise correlations for background power levels (assuming an equal frequency power between all conditions beyond 30Hz), frequency power in the two ranges of interest are higher in the fixation task than in the target detection task (Friedman non-parametric test, all pairwise comparisons, p<0.001), in agreement with the proposal that cognitive flexibility coincides with lower amplitude beta oscillations^52^ and that attentional engagement coincides with lower amplitude alpha oscillations^53, 54^. Importantly, these oscillations are absent from the raw MUA signals (Friedman non-parametric test, all pairwise comparisons, p>0.2), as well as when noise correlations are computed during the same task epochs but from neuronal activities aligned onto target presentation (or saccade go signal in the memory guided saccade task, Friedman non-parametric test, all pairwise comparisons, p>0.2).

Consistent with our previous study ^24^, in all of the three tasks, behavioral performance, defined as the proportion of correct trials as compared to error trials, varied as a function of alpha and beta noise correlation oscillations. Indeed, on a session by session basis, we could identify an optimal alpha (10-16Hz) phase for which the behavioral performance was maximized, in antiphase with a bad alpha phase, for which the behavioral performance was lowest (figure 5C). These effects were highest in the fixation task (34.6% variation in behavioral performance) and lowest though significant in the memory-guided saccade task (13.3% in the target detection task and 9.5% in the memory guided saccade task). Similarly, an optimal beta (20-30Hz) phase was also found to modulate behavioral performance in the same range as the observed alpha behavioral modulations (28.3% variation in behavioral performance in the fixation task, 19.2% in the target detection task and 11% in the memory guided saccade task). As a result, Alpha and beta oscillation phase in noise correlations were predictive of behavioral performance, and the strength of these effects co-varied with alpha and beta oscillation amplitude in noise correlations, being higher in the fixation task, than in the target detection task than in the memory guided saccade task.

### Spike-LFP phase coupling (SFC) varies as a function of task demand

In an independent study^24^, we demonstrate that oscillations in noise correlations arise from specific phase coupling mechanisms between long-range incoming LFP signals and local spiking mechanisms, independently from phase-amplitude coupling mechanisms. Figure 5D represents SFC between spiking activity and LFP signals (see Materials and Methods) computed during a 1200ms time interval starting 300ms after either fixation onset (Fixation and Target detection task) or cue offset (Memory guided saccade task). SFC peaks at both the frequency ranges identified in the noise correlation spectra, namely the high alpha range (10-16Hz) and the beta range (20-30Hz). Importantly, this SFC modulation is highest for the fixation task as compared to the target detection task, thus matching the oscillatory power differences observed in the noise correlations. SFC are lowest in the memory guided saccade task whether considering preferred or non-preferred spatial processing. This is probably due to the fact that the cue to go signal interval of the memory guided saccade task involves memory processes that are expected to desynchronize spiking activity with respect to the LFP frequencies of interest ^46^. This will need to be further explored.

In figure 3, we show layer specific effects onto noise correlations that build up onto the global task effects. An important question is whether these layer effects result from layer specific changes in SFC. Figure 5E represents the SFC data of figure 5D, segregated on the bases of the attribution of the MUA to either superficial or deep cortical FEF layers. While SFC modulations are observed in the same frequencies of interest as in figure 5D, clear layer specific differences can be observed (figure 5F). Specifically, beta ranges SFC are markedly significantly lower in the superficial layers than in the deep layers, for both the detection task and the memory guided saccade task. This points towards a selective control of correlated noise in input, superficial FEF layers. In contrast, alpha range SFC are significantly lower in the superficial layers than in the deep layers only in the memory guided saccade, and specifically when spatial memory is oriented towards a non-preferred location. This points towards overall weaker layer differences for alpha SFC. Alternatively, alpha SFC could result from a different mechanism than beta SFC. This will need to be further explored.

In figure 4 we show that noise correlations dynamically adjust to the probabilistic structure of the trial, and are lowest at the time of highest behavioral demand in the trial. In figure 5D-F, as well as in our independent study^24^, noise correlations and SFC show a strong pattern of co-variation. We propose that noise correlations arise from specific changes in SFC coupling. If this is indeed the case, a strong prediction is SFC will also vary as a function of the probabilistic structure of the task. Figure 6 represents spikes-LFP phase coupling, for each task, for alpha (upper histogram) and beta (lower histogram) frequency ranges during trials with both low and high expected response probability. Importantly, only beta ranges SFC are significantly higher within trials with high expected response probability than in trials with low expected response probability, and this for all tasks.

**Figure 6:**
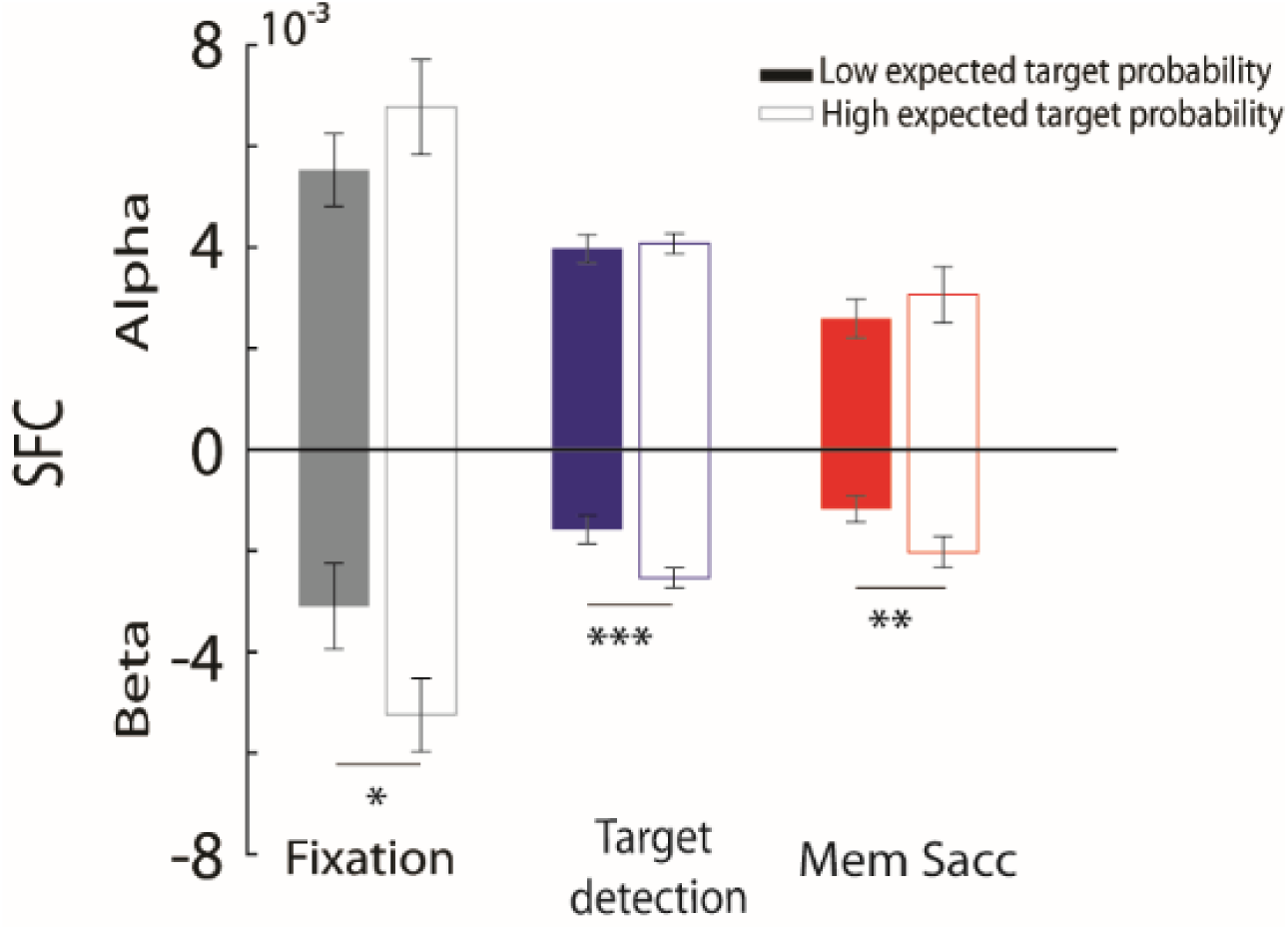
Spike-LFP phase coupling as a function of expected response probability. Average SFC (+/- s.e.) in alpha (10-16Hz, top histogram) and beta (20-30Hz, bottom histogram) for each task and for both trials with low (filled bars) and high (empty bars) expected response probability (t-test, ***: p<0.001).

## Discussion

In this work, our main goal was to examine the impact of cognitive engagement and task demands onto the neuronal population shared variability as assessed from inter-neuronal noise correlations at multiple time scales. Recordings were performed in the macaque frontal eye fields, a cortical region in which neuronal noise correlations have been shown to vary as a function of spatial attention^25^ and spatial memory^26, 55^. Noise correlations were computed over equivalent behavioral task epochs, prior to response production, during a delay in which eyes were fixed and in the absence of any intervening sensory event or motor response. As a result, any observed differences in noise correlations are to be assigned to an endogenous source of shared neuronal variability.

Overall, we demonstrate, for the first time, that noise correlations dynamically adjust to task demands at different time scales. Specifically, we show that noise correlations decrease as cognitive engagement and task demands increase. These task-related variations in noise correlations co-exist with within-trial dynamic changes related to the probabilistic structure of the tasks as well as with long- and short-range oscillatory brain mechanisms.

### Shared neuronal population response variability dynamically adjusts to the behavioral demands

Noise correlations have been shown to vary with learning or changes in behavioral state (V1^56–59^, V4^25, 60–62^ and MT^4, 63, 64^). For example, shared neuronal population response variability was lower in V1 in trained than in naïve monkeys^27^. More recently, Ni et al. describe, within visual areas, a robust relationship between correlated variability and perceptual performance, whether changes in performance happened rapidly (attention instructed by a spatial cue) or slowly (learning). This relationship was robust even when the main effects of attention and learning were accounted for^28^. Here, we question whether changes in noise correlations can be observed simultaneously at multiple time scales. We describe two different times scales at which noise correlations dynamically adjust to the task demands.

The first adjustment in noise correlations we describe is between tasks, that is between blocked contexts of varying cognitive demand, the monkeys knowing that general task requirements will be constant over a hundred of trials or more. Task performance is taken as a proxy to cognitive adjustment to the task demands and negatively correlates with noise correlations in the recorded population. Shared neuronal population variability measure is largest in the fixation task as compared to the two other tasks, by almost 30%. The difference between noise correlations in the target detection task as compared to the guided memory saccade task is in the range of 2%, closer to what has been previously reported in the context of noise correlation changes under spatial attention^25^ or spatial memory manipulations. Importantly, these changes in noise correlations are observed in the absence of significant variations in individual neuronal spiking statistics (average spiking rates, spiking variability or associated Fano factor). To our knowledge, this is the first time that such task effects are described onto noise correlations. This variation in noise correlations as a function of cognitive engagement and task requirements suggests an adaptive mechanism that adjusts noise correlations to the ongoing behavior. Such a mechanism is expected to express itself at different timescales, ranging from the task level, to the across trial level to the within trial level. This is explored next.

It is unclear whether the transitions between high and low noise correlation states when changing from one task to another are fast (over one or two trials) or slow (over tens of trials). In^51^, we show that noise correlations vary as a function of immediate trial past history. Specifically, noise correlations are significantly higher on error trials than on correct trials, both measures being higher if the previous trial is an error trial than if the previous trial is a correct trial. We thus predict a similar past history effect to be observed on noise correlations at transitions between tasks, and we expect for example, noise correlations to be lower in fixation trials that are preceded by a target detection trial, than in trials preceded by fixation trials. In our experimental design, task transitions are unfortunately rare events, precluding the computation of noise correlations on these transitions.

However, our experimental design affords an analysis at a much finer timescale, i.e. the description of a dynamical adjustment in noise correlations within trials. Specifically, we show that noise correlations dynamically adjust to the probability of occurrence of a behaviorally key task event associated with the reward response production (target presentation on the fixation and target detection tasks or saccade go signal on the memory guided saccade task). In other words, shared neuronal population response variability dynamically adjusts to higher demand task epochs. As expected from the general idea that low noise correlations allow for optimal signal processing^12, 65, 66^, we show that, on each of the three tasks, at any given time in the fixation epoch prior to response production, the higher the probability of having to initiate a response, the lower the noise correlations.

Overall, this supports the idea that noise correlations is a flexible physiological parameter that dynamically adjusts at multiple timescales to optimally meet ongoing behavioral demands, as has been demonstrated in multisensory integration^67^ and through learning and attention^28^. The mechanisms through which this possibly takes place are discussed below.

### Rhythmic fluctuations in noise correlations

In the above, we describe changes in noise correlations between tasks as a function of the cognitive demand, as well as within trials, as a function of the probabilistic structure of each task. In addition to these task-related dynamics, we also confirm our concurrent observation of rhythmic fluctuations in noise correlations^24^. These fluctuations are clearly identified in the high alpha frequency range (10-16 Hz) and to a lesser extent in the low gamma frequency range (20-30Hz). To our knowledge, this is the first time that such rhythmic variations in noise correlations are reported. The question is whether these oscillations have a functional relevance or not.

From a behavioral point of view, we show that overt behavioral performance in the three tasks co-varies with both the 10-16Hz and 20-30Hz noise correlation oscillations. In other words, these oscillations account for more than 10% of the behavioral response variability, strongly supporting a functional role for these alpha and beta oscillations.

From a functional point of view, attention directed to the receptive field of neurons has been shown to both reduce noise correlations^24^ and spike-field coherence in the gamma range (V4^68^, it is however to be noted that Engel et al. describe increased spike-field coherence in V1, the gamma range under the same conditions, hinting towards areal specific differences^69^). In our hands, both the rhythmic fluctuations in noise correlations and the task and trial-related changes in noise correlations co-exist with increased spike-LFP phase coupling in the very same 10-16Hz and/or 20-30Hz frequency ranges we identify in the noise correlations. This suggests that changes in shared neuronal variability possibly arise from changes in the local coupling between neuronal spiking activity and local field potentials. Supporting such a functional coupling, both the observed changes in noise correlations and spike-LFP phase coupling in the frequencies of interest are highest in the fixation task as compared to the other two tasks.

Beta oscillations in the local field potentials (LFP) are considered to reflect long-range processes and have been associated with cognitive control and flexibility^29, 52, 69–72^ as well as with motor control^73–75^(for review see^52^). Specifically, lower beta power LFPs has been associated with states of higher cognitive flexibility. In our hands, lower beta in noise correlations correspond to higher cognitive demands. We thus hypothesize a functional link between these two measures, LFP oscillations locally changing spiking statistics, i.e. noise correlations, by a specific spike-LFP phase coupling in this frequency range. Supporting a long-range origin of these local processes (figure 7, inset), we show that spike-LFP phase coupling in this beta range strongly decreases in the more superficial cortical layers as compared to the deeper layers, as task cognitive demand increases. On the other hand, alpha oscillations are associated with attention, anticipation^53, 54^, perception^76–78^, and working memory^79^. As for beta oscillations, lower alpha in noise correlations, and accordingly in spike-LFP phase coupling, correspond to higher cognitive demands. In contrast with what is observed for beta spike-LFP phase coupling, alpha spike-LFP phase coupling does not exhibit any layer specificity across task demands. Thus overall, alpha and beta rhythmicity account for strong fluctuations in behavioral performance, as well as for changes in spike-LFP phase coupling. However, beta processes seem to play a distinct functional role as compared to the alpha processes, as their effect is more marked in the superficial than in the deeper cortical layers. These observations coincide with recent evidence that cognition is rhythmic^80, 81, 87^ and that noise correlations play a key role in optimizing behavior to the ongoing time-varying cognitive demands^27^.

## Acknowledgments

S.B.H.H. was supported by ANR grant ANR-14-CE13-0005-1. C.G. was supported by the French Ministère de l’Enseignement Supérieur et de la Recherche. S.B.H. was supported by ANR grant ANR-11-BSV4-0011, ANR grant ANR-14-CE13-0005-1, and the LABEX CORTEX (ANR-11-LABX-0042) of Université de Lyon, within the program Investissements d’Avenir (ANR-11-IDEX-0007) operated by the French National Research Agency (ANR). E.A. was supported by the CNRS-DGA and Fondation pour la Recherche Médicale. We thank research engineer Serge Pinède for technical support and Jean-Luc Charieau and Fabrice Hérant for animal care. All procedures were approved by the local animal care committee (C2EA42-13-02-0401-01) in compliance with the European Community Council, Directive 2010/63/UE on Animal Care.

## Authors contributions

Conceptualization, S.B.H. S.B.H.H. and C.G.; Methodology, S.B.H., S.B.H.H., C.G., E.A., C.W.; Investigation, S.B.H., S.B.H.H., C.G., E.A. and C.W.; Writing – Original Draft, S.B.H. S.B.H.H. and C.G.; Writing – Review & Editing, S.B.H. S.B.H.H. and C.G.; Funding Acquisition, S.B.H.; Supervision, S.B.H.

## Supplementary material and methods

### Material and methods

#### Ethical statement

All procedures were in compliance with the guidelines of European Community on animal care (Directive 2010/63/UE of the European Parliament and the Council of 22 September 2010 on the protection of animals used for scientific purposes) and authorized by the French Committee on the Ethics of Experiments in Animals (C2EA) CELYNE registered at the national level as C2EA number 42 (protocole C2EA42-13-02-0401-01).

#### Surgical procedure

As in^51^, two male rhesus monkeys (Macaca mulatta) weighing between 6-8 kg underwent a unique surgery during which they were implanted with two MRI compatible PEEK recording chambers placed over the left and the right FEF hemispheres respectively (figure 1A), as well as a head fixation post. Gas anesthesia was carried out using Vet-Flurane, 0.5 – 2% (Isofluranum 100%) following an induction with Zolétil 100 (Tiletamine at 50mg/ml, 15mg/kg and Zolazepam, at 50mg/ml, 15mg/kg). Post-surgery pain was controlled with a morphine pain-killer (Buprecare, buprenorphine at 0.3mg/ml, 0.01mg/kg), 3 injections at 6 hours interval (first injection at the beginning of the surgery) and a full antibiotic coverage was provided with Baytril 5% (a long action large spectrum antibiotic, Enrofloxacin 0.5mg/ml) at 2.5mg/kg, one injection during the surgery and thereafter one each day during 10 days. A 0.6mm isomorphic anatomical MRI scan was acquired post surgically on a 1.5T Siemens Sonata MRI scanner, while a high-contrast oil filled grid (mesh of holes at a resolution of 1mm×1mm) was placed in each recording chamber, in the same orientation as the final recording grid. This allowed a precise localization of the arcuate sulcus and surrounding gray matter underneath each of the recording chambers. The FEF was defined as the anterior bank of the arcuate sulcus and we specifically targeted those sites in which a significant visual and/or oculomotor activity was observed during a memory guided saccade task at 10 to 15° of eccentricity from the fixation point (figure 1A). In order to maximize task-related neuronal information at each of the 24-contacts of the recording probes, we only recorded from sites with task-related activity observed continuously over at least 3 mm of depth.

#### Behavioral task

During a given experimental session, the monkeys were placed in front of a computer screen (1920×1200 pixels and a refresh rate of 60 Hz) with their head fixed. Their water intake was controlled so that their initial daily intake was covered by their performance in the task, on a trial by trial basis. This quantity was complemented as follows. On good performance sessions, monkeys received fruit and water complements. On bad performance sessions, water complements were provided at a distance from the end of the session. Each recording session consisted of random alternations of three different tasks (see below and figure 1B), so as to control for possible time in the session or task order effects. For all tasks, to initiate a trial, the monkeys had to hold a bar in front of the animal chair, thus interrupting an infrared beam. (1) ***Fixation Task*** (figure 1B.1): A red fixation cross (0.7×0.7°), appeared in the center of the screen and the monkeys were required to hold fixation during a variable interval randomly ranging between 7000 and 9500ms, within a fixation window of 1.5×1.5°, until the color change of the central cross. At this time, the monkeys had to release the bar within 150-800 ms after color change. Success conditioned reward delivery. (2) ***Target detection Task*** (figure 1B.2): A red fixation cross (0.7×0.7°), appeared in the center of the screen and the monkeys were required to hold fixation during a variable interval ranging between 1300 and 3400 ms, within a fixation window of 1.5×1.5°, until a green squared target (0.28×0.28°) was presented for 100 ms in one of four possible positions ((10°,10°), (−10°,10°), (−10°,−10°) and (10°,−10°)) in a randomly interleaved order. At this time, the monkeys had to release the bar within 150-800 ms after target onset. Success conditioned reward delivery. (3) ***Memory-guided saccade Task*** (figure 1B.3): A red fixation cross (0.7×0.7°) appeared in the center of the screen and the monkeys were required to hold fixation for 500 msec, within a fixation window of 1.5×1.5°. A squared green cue (0.28×0.28°) was then flashed for 100ms at one of four possible locations ((10°, 10°), (−10°, 10°), (−10°,−10°) and (10°,−10°)). The monkeys had to continue maintain fixation on the central fixation point for another 700-1900 ms until the fixation point disappeared. The monkeys were then required to make a saccade towards the memorized location of the cue within 500-800ms from fixation point disappearance, and a spatial tolerance of 4°x4°. On success, a target, identical to the cue was presented at the cued location and the monkeys were required to fixate it and detect a change in its color by a bar release within 150-800 ms from color change. Success in all of these successive requirements conditioned reward delivery.

#### Neural recordings

On each session, bilateral simultaneous recordings in the two FEFs were carried out using two 24-contacts Plexon U-probes. The contacts had an interspacing distance of 250 µm. Neural data was acquired with the Plexon Omniplex® neuronal data acquisition system. The data was amplified 400 times and digitized at 40,000 Hz. The MUA neuronal data was high-pass filtered at 300 Hz. The LFP neuronal data was filtered between 0.5 and 300 Hz. In the present paper, all analyses are performed on the multi-unit activity recorded on each of the 48 recording contacts. A threshold defining the multi-unit activity was applied independently for each recording contact and before the actual task-related recordings started. All further analyses of the data were performed in Matlab™ and using FieldTrip ^82^ and the Wavelet Coherence Matlab Toolbox ^83^, both open source Matlab™ toolboxes.

### Data Analysis

#### Data preprocessing

Overall, MUA recordings were collected from 48 recording channels on 26 independent recording sessions (13 for M1 and 13 for M2). We excluded from subsequent analyses all channels with less than 5 spikes per seconds. For each session, we identified the task-related channels based on a statistical change (one-way ANOVA, p<0.05) in the MUA neuronal activity in the memory-guided saccade task, in response to either cue presentation ([0 400] ms after cue onset) against a pre-cue baseline ([-100 0] ms relative to cue onset), or to saccade execution go signal and to saccade execution (i.e. fixation point off, [0 400] ms after go signal) against a pre-go signal baseline ([-100 0] ms relative to go signal), irrespective of the spatial configuration of the trial. In total, 671 channels were retained for further analyses out of 1248 channels.

#### MUA spatial selectivity

FEF neurons are characterized by a strong visual, saccadic, spatial memory and spatial attention selectivity^30, 38, 39^. We used a one-way ANOVA (p<0.05) to identify the spatially selective channels in response to cue presentation ([0 400] ms following cue onset) and to the saccade execution go signal ([0 400] ms following go signal). Post-hoc t-tests served to further order, for each channels, the neuron’s response in each visual quadrant from preferred (p1), to least preferred (p4). By convention, positive modulations were considered as preferred and negative modulations as least preferred. For example, in a given session, the MUA signal recorded on channel 1 of a probe placed in the left FEF, could have as best preferred position p1 the upper right quadrant, the next best preferred position p2 the lower right quadrant, the next preferred position p3 the upper left quadrant and the least preferred position p4 the lower left quadrant. The MUA signal recorded on channel 14 of this same probe, could have as best preferred position p1 the lower right quadrant, the next best preferred position p2 the upper right quadrant, the next preferred position p3 the lower left quadrant and the least preferred position p4 the upper left quadrant. Positions with no significant modulation in any task epoch were labeled as p0 (no selectivity for this position). Once this was done, for each electrode, pairs of MUA recordings were classified along two possible functional categories: pairs with the same spatial selectivity (SSS pairs, sharing the same p1) and pairs with different spatial selectivities (DSS pairs, such that the p1 of one MUA is a p0 for the other MUA). For the sake of clarity, we do not consider partial spatial selectivity pairs (such that the p1 of one MUA is a non-preferred, p2, p3 or p4 for the other MUA).

#### MUA layer attribution

As stated above, our recordings are not tangential to cortical surface. As a proxy to attribute a given recording channel to upper or lower cortical layers we proceeded as follows. For each electrode and each channel, we estimated, at the time of cue onset in the memory-guided saccade task (100ms-500ms from cue onset), the spike-field coherence in the alpha range (6 to 16 Hz) and the gamma range (40 to 60 Hz). Based on previous literature ^84^, we used the ratio between the alpha and gamma spike field-coherence as a proxy to assign the considered LFP signals to a deep cortical layer site (high alpha / gamma spike-field coherence ratio) or to a superficial cortical layer site (low alpha / gamma spike-field coherence ratio). We also categorized MUA signals into visual, visuo-motor and motor categories, as in Cohen et al. (2009). Briefly, average firing rates were computed in 3 epochs: [-100 0] ms before cue onset (baseline), [0 200] ms after cue onset (visual), and [0 200] ms before saccade onset (movement). Neurons with activity statistically significantly different from the baseline (Wilcoxon rank-sum test, *P* < 0.05) after cue onset were categorized as visual. Neurons with activity statistically significantly different from the baseline (Wilcoxon rank-sum test, *P* < 0.05) before saccade onset were categorized as oculomotor. Neurons that were active in both epochs were categorized as visuo-movement neurons. The LFP categorization along the alpha to gamma spike-field coherence ratio strongly coincided with the classification of the MUA signals into purely visual sites (low alpha and gamma spike-field coherence ratio, input FEF layers) and visuo-motor sites (high alpha and gamma spike-field coherence ratio, output FEF layers, figure 4).

#### Noise Correlations

The aim of the present work is to quantify task effects onto the spiking statistics of the FEF spiking activity during equivalent task-fixation epochs. The statistics that we discuss is that of noise correlations between the MUA activities on the different simultaneously recorded signals. For each channel, and each task, intervals of interest of 200ms were defined during the fixation epoch from 300 ms to 500 ms from eye fixation onset. Specifically, for each channel i, and each trial k, the average neuronal response r_i_(k) for this time interval was calculated and z-score normalized into z_i_(k), where z_i_(k)=r_i_(k)-µ_i_/stdi and µ_i_ and std_i_ respectively correspond to the mean firing rate and standard deviation around this mean during the interval of interest of the channel of interest i. This z-score normalization allows to capture the changes in neuronal response variability independently of changes in mean firing rates. Noise correlations between pairs of MUA signals during the interval of interest were then defined as the Pearson correlation coefficient between the z-scored individual trial neuronal responses of each MUA signal over all trials. Only positive significant noise correlations are considered, unless stated otherwise. In any given recording session, noise correlations were calculated between MUA signals recorded from the same electrode, thus specifically targeting intra-cortical correlations. This procedure was applied independently for each task. Depending on the question being asked, noise correlations were either computed on activities aligned on fixation onset, or on activities aligned on target (Fixation and Target detection task) or saccade execution (memory guided saccade task) signals.

In order to control for the fact that the observed changes in noise correlations cannot be attributed to changes in other firing rate metrics, several statistics were also extracted, from comparable task epochs, from 300 to 500ms following trial initiation and fixation onset. None of these metrics were significantly affected by the task. Specifically, we analyzed (a) mean firing rate (ANOVA, p>0.5), (b) the standard error around this mean firing rate (ANOVA, p>0.6), and (c) the corresponding Fano factor (ANOVA, p>0.7). These data, reproducing previous reports ^51, 85^ are not shown.

#### Oscillations in noise correlations

To measure oscillatory patterns in the noise correlation time-series data, we computed, for each task, and each session (N=12), noise correlations over time (over successive 200ms intervals, sliding by 10ms, running from 300ms to 1500ms following eye fixation onset for Fixation and Target detection tasks and from 300ms to 1500ms following cue offset form Memory-guided saccade task). A wavelet transform ^82^ was then applied on each session’s noise correlation time series. Statistical differences in the noise correlation power frequency spectra were assessed using a non-parametric Friedman test. When computing the noise correlations in time, we equalized the number of trials for all tasks and all conditions so as to prevent any bias that could be introduced by unequal numbers of trials. To control that oscillations in noise correlations in time cannot be attributed to changes in spiking activity, a wavelet analysis was also run onto MUA time series data (data not shown).

#### Modulation of behavioral performance by phase of noise correlation alpha and beta rhythmicity

To quantify the effect of noise correlation oscillations onto behavioral performance, we used a complex wavelet transform analysis (Fieldtrip, Oostenveld et al. 2011) to compute, for each session and each task, in the noise correlations, the phase of the frequencies of interest (alpha / beta) following eye fixation onset (for the Fixation and Target detection tasks) or cue offset (for the Memory guided saccade task). For each session, we identified hit and miss trials falling at zero phase of the frequency of interest (+/- π /140) with respect to target presentation or fixation point offset time. In the fixation task, premature fixation aborts by anticipatory manual response or eye fixation failure were considered as misses. Hit rates (HR) were computed for this zero phase bin. We then shifted this phase window by π /70 steps and recalculated the HR, repeating this procedure to generate phase-detection HR functions, across all phases, for each frequency of interest ^86^. For each session, the phase bin for which hit rate was maximal was considered as the optimal phase. The effect of a given frequency (alpha or beta) onto behavior corresponds to the difference between HR at this optimal phase and HR at the anti-optimal phase (optimal phase + π). To test for statistical significance, observed hit/miss phases were randomized across trials so as to shuffle the temporal relationship between phases and behavioral performance. This procedure was repeated 1000 times. 95% CI was then computed and compared to the observed behavioral data.

#### Spikes-LFP Phase coupling

For each selected channel, spikes-LFP phase coupling spectra (SFC) were calculated between the spiking activity obtained in one channel and the LFP activity from the next adjacent channel in the time interval running from 300ms to 1500ms following cue offset. We used a single Hanning taper and applied convolution transform to the Hanning-tapered trials. We equalized the number of trials for all conditions so as to prevent any bias that could be introduced by unequal numbers of trials. We used a 4 cycles length per frequency. The memory guided saccade task is known to involve spatial processes during the cue to target interval that bias spike field coherence. Thus, spikes-LFP phase coupling was measured separately for trials in which the cued location matched the preferred spatial location of the channel and trials in which the cued location did not match the preferred spatial location of the channel. Statistics were computed across channels x sessions, using a non-parametric Friedman test.

## Supplementary figures

**Figure S1:**
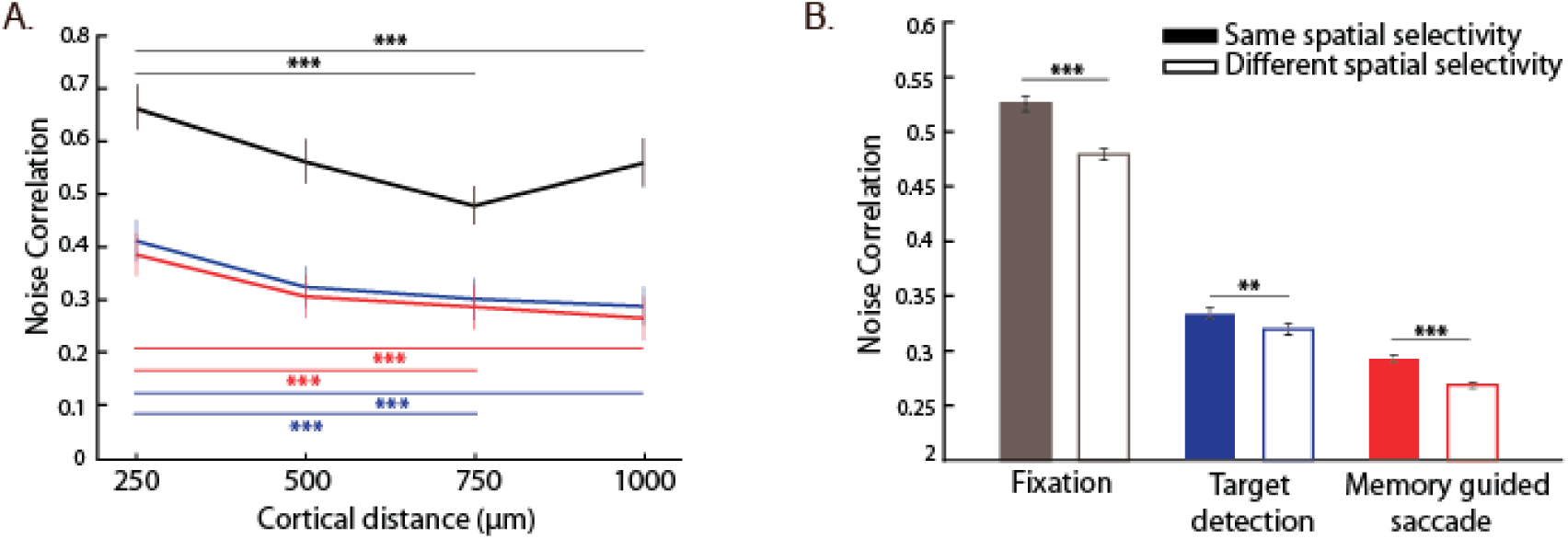
(A) ***Noise correlations as a function of cortical distance*.** Average noise correlations (mean +/- s.e.) across sessions, for each task (conventions as in (A)), from 300 ms to 500ms after eye fixation onset, as a function of distance between pairs of channels: 250µm; 500µm; 750µm; 1000µm. Stars indicate statistical significance following a two-way ANOVA and ranksum post-hoc tests; *p<0.05; **p<0.01; ***p<0.001. (B) ***Noise correlations as a function of spatial selectivity***. Average noise correlations (mean +/- s.e.) across sessions, for each tasks (conventions as in figure 2), from 300ms to 500ms after eye fixation onset, as a function of whether noise correlations are calculated over signals sharing the same spatial selectivity (full bars) or not (empty bars). Stars indicate statistical significance following a two-way ANOVA and ranksum post-hoc tests; *p<0.05; **p<0.01; ***p<0.001.

